# Erythritol synthesis in human cells is elevated in response to oxidative stress and regulated by the non-oxidative pentose phosphate pathway

**DOI:** 10.1101/2022.03.07.483290

**Authors:** Semira R. Ortiz, Alexander Heinz, Karsten Hiller, Martha S. Field

**Affiliations:** Division of Nutritional Sciences, Cornell University, Ithaca, NY 14853, USA; Department of Bioinformatics and Biochemistry, BRICS, Technische Universität Braunschweig, 38106 Braunschweig, Germany

**Keywords:** erythritol, oxidative stress, pentose phosphate pathway

## Abstract

Erythritol is a predictive biomarker of cardiometabolic diseases and is produced from glucose metabolism through the pentose phosphate pathway (PPP). Little is known regarding the regulation of endogenous erythritol synthesis in humans. In the present study, we investigated the stimuli that promote erythritol synthesis in human cells and characterized potential points of regulation along the PPP. Human A549 lung carcinoma cells were chosen for their known ability to synthesize erythritol. A549 cells were treated with potential substrates for erythritol production, including glucose, fructose, and glycerol. Using siRNA knockdown, we assessed the necessity of enzymes G6PD, TKT, TALDO, and SORD for erythritol synthesis. We also used position-specific ^13^C-glucose tracers to determine whether the carbons for erythritol synthesis are derived directly from glycolysis or through the oxidative PPP. Finally, we assessed if erythritol synthesis responds to oxidative stress using chemical and genetic models. Intracellular erythritol was directly associated with media glucose concentration. In addition, siRNA knockdown of TKT or SORD inhibited erythritol synthesis, whereas siG6PD did not. Both chemically induced oxidative stress and constitutive activation of the antioxidant response transcription factor NRF2 elevated intracellular erythritol. Our findings indicate that erythritol synthesis is proportional to flux through the PPP and is regulated by non-oxidative PPP enzymes.

## 1. Introduction

Serum erythritol is a predictive biomarker of chronic disease onset and associated complications. In one large prospective cohort, baseline serum erythritol was elevated in subjects who developed cardiovascular disease (CVD) or type 2 diabetes mellitus (T2DM) up to 20 years later [1,2]. Another recent study compared patients with cardiovascular risk factors who did and did not develop coronary artery disease. Serum erythritol was significantly elevated in those who did develop coronary artery disease. Erythritol has also been shown to predict risk for diabetic complications including retinopathy, nephropathy, and arterial stiffness [3–5]. Serum erythritol appears to be an early, general marker of cardiometabolic dysfunction.

Erythritol is a four-carbon polyol that was recently found to be endogenously synthesized in humans. Little is known regarding the regulation of erythritol synthesis. Hootman et al. first demonstrated that erythritol is produced from glucose in humans through the pentose phosphate pathway (PPP) [6]. It was further identified that the enzymes sorbitol dehydrogenase (SORD) and alcohol dehydrogenase 1 (ADH1) are responsible for catalyzing the final step in erythritol synthesis from glucose (namely conversion of erythrose to erythritol) using the cofactor NADPH [7].

The PPP branches off from glycolysis and fuels anabolic reactions. It consists of two phases: the oxidative PPP and the non-oxidative PPP. The oxidative PPP generates NADPH, which is essential for endogenous antioxidant generation and lipid synthesis. The non-oxidative PPP consists of a series of sugar interconversions that provide precursors for nucleotide synthesis. The non-oxidative phase can convert pentoses back to glycolytic intermediates through reversible reactions. PPP metabolism plays roles in the development of cardiometabolic diseases across multiple organs. The rate limiting enzyme of the PPP, glucose-6-phosphate dehydrogenase (G6PD), is elevated in the adipose tissue and skeletal muscle of prediabetic subjects [8,9]. In adipose tissue, elevated G6PD promotes pro-inflammatory macrophages, exacerbating insulin resistance [9]. In skeletal muscle cells, inhibition of G6PD can improve insulin-stimulated glucose uptake [8]. PPP flux is also important in the liver, where NADPH synthesis can contribute to fatty liver development [10].

As a product of the PPP, erythritol synthesis may be an indicator of high PPP flux and the associated aberrant changes in glucose metabolism. The purpose of this study was to identify the upstream factors that regulate erythritol synthesis. Our findings highlight that erythritol synthesis is increased in response to PPP stressors and is primarily regulated by glucose availability and activity of enzymes within the non-oxidative PPP.

## 2. Methods

### 2.1 Cell culture and treatment with carbohydrates

A549 cells were obtained from ATCC (CCL-185) and maintained in Minimum Essential Medium Alpha Modification (MEM Alpha) containing ribonucleosides, deoxyribonucleosides, phenol red, and l-glutamine, 1% penicillin/streptomycin (Cytiva), and 10% FBS (Cytiva). KEAP1 variant cells were maintained in MEM Alpha with the addition of 1 µg/mL puromycin. HK-2 cells were obtained from ATCC (CRL-2190) and cultured in Dulbecco’s Modified Eagle Medium (DMEM) (Corning) with 1% penicillin/streptomycin and 10% FBS (Cytiva).

Unless otherwise noted, cells were seeded at a density of 1.5-2 × 10^5^ cells per well in 6-well plates and allowed to proliferate overnight before treatment. For initial characterization of erythritol production from glucose and fructose, modified glucose-free DMEM (Hyclone) was prepared to contain 6.25, 12.5, or 25 mM glucose or 25 mM fructose. All further measurements (including knockdowns and metabolic assays) were performed in standard MEM Alpha or DMEM (5.5 mM glucose) with additional glucose, mannitol, or glycerol added to achieve desired concentrations of each carbohydrate. Cells were incubated with carbohydrates for 48 h, then polar metabolites were extracted, and relative cell number was measured by MTT.

### 2.2 Extraction and measurement of polar metabolites by GC-MS

Polar metabolites were extracted following the protocol for adherent cells by Sapcariu et al [11]. 10 µM ^13^C1-ribitol (Cambridge Isotope Laboratories) was added to methanol as an internal standard during extraction. Dried extracts were derivatized and metabolites (erythritol, ^13^C1-ribitol, and sorbitol) were measured by GC-MS as previously described [12]. In SIM mode, mass spectra of erythritol (*m/z* 217), ^13^C1-ribitol (*m/z* 218), and sorbitol (*m/z* 319) were acquired from 8-9 min, 10-11 min, and 12-13 min, respectively. Metabolite peaks were selected based on the retention time of their respective standards. Relative erythritol and sorbitol were calculated by dividing their absolute intensity by the absolute intensity of ^13^C1-ribitol. Relative erythritol and sorbitol were then normalized to cell number, measured by MTT.

### 2.3 MTT assay for relative cell number

MTT reagent (MP Bio) dissolved in 1X PBS (5 mg/mL) was added to culture medium to a final concentration of 0.16 mg/mL. Cells were incubated for 4 hours at 37°C, after which media and MTT reagent were removed. Formazan crystals were solubilized in 1 mL DMSO, diluted 1:10 in additional DMSO, then transferred to a microplate to quantify A570 using a Biotek plate reader.

### 2.4 Knockdown of *SORD, G6PD, TKT*, and *TALDO* using siRNA

Non-targeting control siRNA and siRNA targeting *SORD, G6PD, TKT*, and *TALDO* were purchased from Horizon Discovery. Product numbers and sequences are provided in Table S1. A549 cells were reverse transfected using RNAiMAX (Thermo Fisher Scientific) according to manufacturer’s instructions. Per well (6-well plate), 2 µL of 20 µM siRNA and 5 µL RNAiMAX were diluted in 400 µL OptiMEM (Thermo Fisher Scientific), mixed gently and incubated for 20 min at room temperature. The solution was applied to the well 5 min prior cell seeding. 2 × 10^5^ cells were seeded in 1600 µL standard 5.5 mM glucose MEM Alpha or 25 mM glucose MEM Alpha. Transfected cells were incubated for 48 h until metabolite extraction or measurement of cell density by MTT.

### 2.5 Measurement of intracellular NADPH and metabolic phenotype

For measurement of NADPH or metabolic phenotype in 96-well plates, cells were transfected as described with reagents adjusted to a provide final volume of 200 µL per well: 0.1 µL siRNA, 0.25 µL RNAiMAX, and 20 µL OptiMEM, and 180 µL of culture medium.

NADPH was measured following reverse transfection using the NADP/NADPH-Glo™ Assay (Promega). Briefly, 7 × 10^3^ cells per well were reverse transfected with siRNA targeting control, *SORD*, or *G6PD* and incubated for 48 h in standard (5.5 mM glucose) or 25 mM glucose MEM Alpha. To measure NADPH individually, culture media was aspirated and replaced with 50 µL 1X PBS and 50 µL of 0.2 N NaOH with 1% DTAB to lyse cells. 50 µL of cell lysate was transferred to a white 96-well luminometer plate then incubated for 15 min at 60C followed by 10 minutes at room temperature. The base was neutralized with 50 µL Trizma/HCL, then NADPH was measured as described in the manufacturer’s protocol.

Oxygen consumption rate (OCR) and extracellular acidification (ECAR) were measured using the Seahorse XF Cell Energy Phenotype Test Kit (Agilent). 12 × 10^3^ cells per well were reverse transfected with siRNA control or targeting *SORD* in a Seahorse XF24 Cell Culture Microplate (Agilent) with MEM Alpha adjusted to 10 mM glucose. 10 mM glucose was chosen to match the Assay Buffer glucose concentration. After 48 h, baseline OCR and ECAR were measured following the manufacturer’s protocol in Seahorse Assay Buffer (phenol red-free DMEM with 1 mM pyruvate, 2 mM glutamine, and 10 mM glucose).

### 2.6 Western blot analysis

Cells were lifted using 0.25% trypsin-EDTA, pelleted, and rinsed once with 1X PBS. Cell pellets were lysed by sonication in lysis buffer containing 15% NaCl, 5 mM EDTA (pH 8), 1% Triton X100, 10 mM Tris-Cl, 5 mM DTT, and 10 μL/mL protease inhibitor cocktail (Sigma Aldrich). Protein concentration was determined with a modified Lowry assay [13]. Equal amounts of protein (15-25 µg) were loaded onto a 10% SDS gel and transferred to a PVDF membrane (MilliporeSigma). Membranes were blocked overnight at 4C in 5% non-fat milk, incubated in primary antibody overnight at 4C, then incubated with secondary antibody for 1 hr at room temperature. Primary antibodies against alpha tubulin (ATUB), KEAP1, GAPDH, G6PD (Cell Signaling Technology), SORD, TALDO, and TKT (ProteinTech) were diluted 1:1000. Secondary anti-rabbit and anti-mouse antibodies were diluted 1:50,000 (ThermoFisher). After antibody incubation, blots were imaged using a Protein Simple FluorChem E with Clarity Western ECL Substrate (Bio-Rad). Band intensity was measured using ImageJ (NIH).

### 2.7 ^13^C-glucose tracing

To measure the incorporation of labelled glucose in endogenous erythritol, modified glucose-free DMEM (Hyclone) was adjusted to 25 mM glucose with either 1-^13^C-glucose or 6-^13^C-glucose (Cambridge Isotope Laboratories). Cells were reverse transfected as described above, incubated with labelled glucose for 48 h, then polar metabolites were harvested and measured by GCMS. The mass spectra were acquired for erythritol from *m/z* 320 (M0) to *m/z* 324 (M4). Mass isotope distribution was calculated as previously described [6].

### 2.8 Treatment with hydrogen peroxide

Cells were seeded at 2 × 10^5^ cells per well in 6-well plates and allowed to proliferate overnight. The following day, cells were treated with water or hydrogen peroxide ranging from 150-600 µM hydrogen peroxide for either 6 or 8 h. After treatment, polar metabolites and relative cell number were measured. The percentage of live cells was determined using trypan blue staining quantified using a TC10 Automated Cell Counter (Bio-Rad). To determine intracellular NADPH and oxidized glutathione (GSSG), 7 × 10^3^ cells/well in a 96-well plate were treated with hydrogen peroxide, then NADPH was measured as described above. GSSG was measured using the GSH/GSSG-Glo assay (Promega) per the manufacturer’s protocol.

### 2.9 Statistical Analysis

All statistical analyses were conducted in GraphPad Prism 9 (GraphPad Software). All data are shown as mean ± SD, and p-values lower than 0.05 were considered statistically significant. Comparisons between two groups were analyzed by unpaired t-test. Comparisons between more than two groups were analyzed by one-way ANOVA followed by Tukey’s multiple comparisons test or two-way ANOVA with Sidak’s or Tukey’s multiple comparisons test.

## 3. Results

### 3.1 Intracellular erythritol increases in response to glucose and fructose in culture medium

A549 cells were used to assess the contribution of substrate availability to erythritol synthesis given their robust PPP activity and known ability to generate erythritol [7]. As expected, intracellular erythritol was significantly higher in A549 cells cultured in 25 mM glucose compared to 6.25 mM glucose (Fig. 1A, p < 0.0001). In the absence of glucose, 25 mM fructose also significantly increased erythritol compared to the 6.25 mM glucose control (Fig. 1A, p < 0.05). Treatment with 25 mM mannitol was performed as a control for the osmotic stress induced by high-glucose (25 mM) media. Mannitol treatment did not significantly increase intracellular erythritol compared to 5.5 mM glucose controls (Fig. 1B). Sorbitol accumulation is one mechanism of responding to hyperosmolarity [14]. Measurement of intracellular sorbitol confirmed the strong induction of osmotic stress in mannitol-treated cells. Mannitol elevated sorbitol nearly 40-fold above 5.5 mM glucose and 5-fold above 25 mM glucose-treated cells (Fig. 1C, p < 0.0001). Sorbitol accumulation was also modestly induced by 25 mM glucose compared to control media, as expected due to increased flux through the polyol pathway as a result of increased glucose (Fig. 1C, p < 0.05)[14].

**Figure 1.**
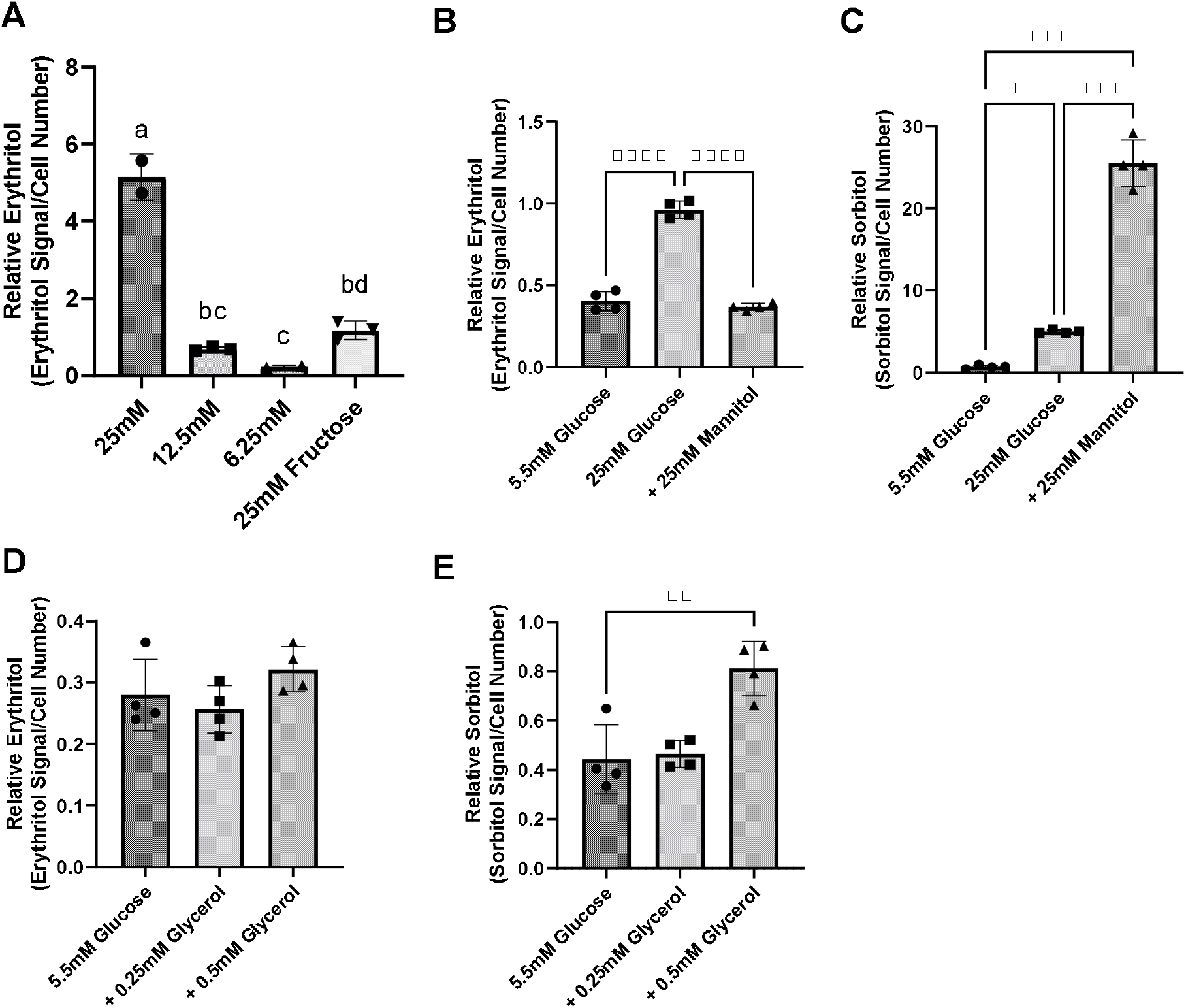
Intracellular erythritol and sorbitol levels respond to carbohydrate concentrations in media and osmotic stress. (A) Relative erythritol in cells cultured for 48 hrs in media containing 25mM, 12.5mM, or 6.25 mM glucose or 25mM fructose. Labelled means without a common letter differ, p < 0.05. (B) Relative erythritol and (C) relative sorbitol in cells cultured in media containing 5.5mM or 25mM glucose, or 5.5mM glucose supplemented with 25mM mannitol. (D) Relative erythritol and (E) relative sorbitol in cells cultured in 5.5mM glucose media or 5.5mM glucose media containing 0.25mM or 0.5mM glycerol. All relative metabolite values are normalized to internal standard and cell number. Data are shown as mean ± SD and were analyzed by one-way ANOVA, followed by Tukey’s multiple comparisons test (n=2-4). *p<0.05, **p<0.01, ****p<0.0001

We also evaluated the effect of exposure to excess glycerol, which acts as both an osmolyte and an alternative carbon source, on erythritol synthesis. There was no significant difference in erythritol content in cells treated with 5.5 mM glucose and 5.5 mM glucose media containing 0.25 mM or 0.5 mM glycerol (Fig. 1D). In contrast, sorbitol was significantly higher in cells treated with 0.5 mM glycerol compared to control cells (Fig. 1E, p < 0.01).

We have previously demonstrated that in mice, the liver and kidney are primary contributors to endogenous erythritol synthesis [12]. We therefore repeated exposure to 5.5 mM or 25 mM glucose media in HK-2 cells, a human proximal tubular cell line derived from normal kidney cells. High glucose media caused a 40% increase in erythritol in HK-2 cells (Fig. 2, p < 0.001). Collectively, these findings demonstrate that erythritol synthesis is elevated in response to glucose availability and not because of osmotic stress.

**Figure 2.**
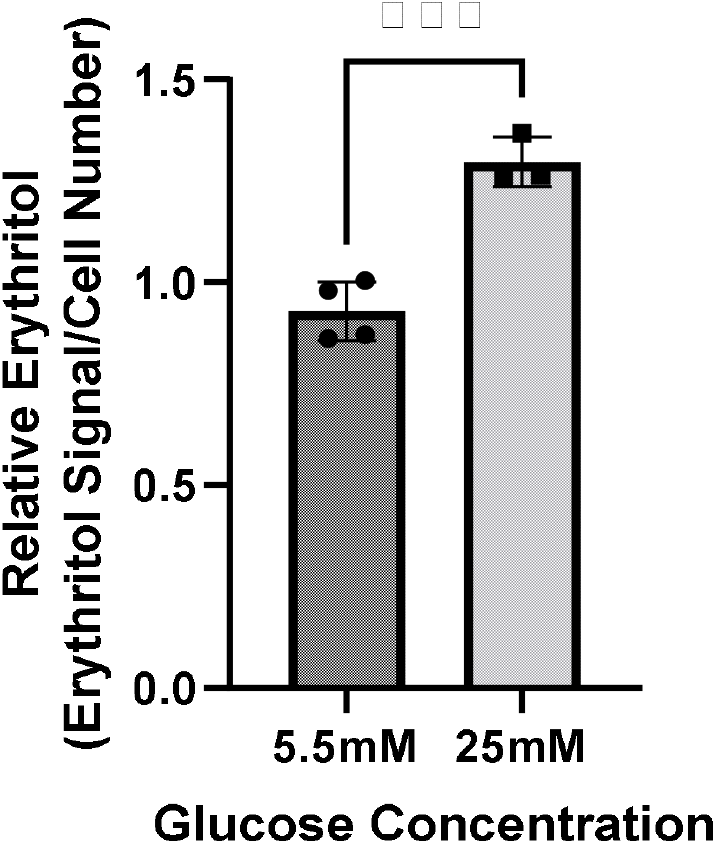
Intracellular erythritol in HK-2 cells is increased in high-glucose media. Intracellular erythritol was measured in HK-2 cells exposed to 5.5mM or 25mM glucose media for 48hrs (n=4). Data is normalized by internal standard and cell number, shown as mean ± SD, and analyzed by unpaired t-test. ***p<0.001.

### 3.2 Reduction of SORD decreases erythritol synthesis and NADPH availability

As previously described, erythritol is synthesized from glucose through the PPP in mammalian cells [6,7]. To determine the rate-limiting step in erythritol synthesis, we reduced expression of SORD and G6PD using siRNA. G6PD is generally considered the rate-limiting enzyme of the PPP. Unexpectedly, we found that in 5.5 mM glucose media, reduction of neither enzyme affected intracellular erythritol levels (Fig. 3A). Knockdown of SORD and G6PD were validated by western blot (Fig. 3B). When exposed to 25 mM glucose media, siControl and siG6PD cells both had significantly higher erythritol compared to cells cultured in 5.5 mM glucose (Fig. 3A, p < 0.001 and p < 0.0001, respectively). siSORD cells did have reduced intracellular erythritol in response to hyperglycemia (Fig. 3A), consistent with previous findings [7]. These data demonstrate that, in A549 cells, erythritol synthesis is limited by *SORD* expression but not by *G6PD* expression in response to high-glucose culture medium.

**Figure 3.**
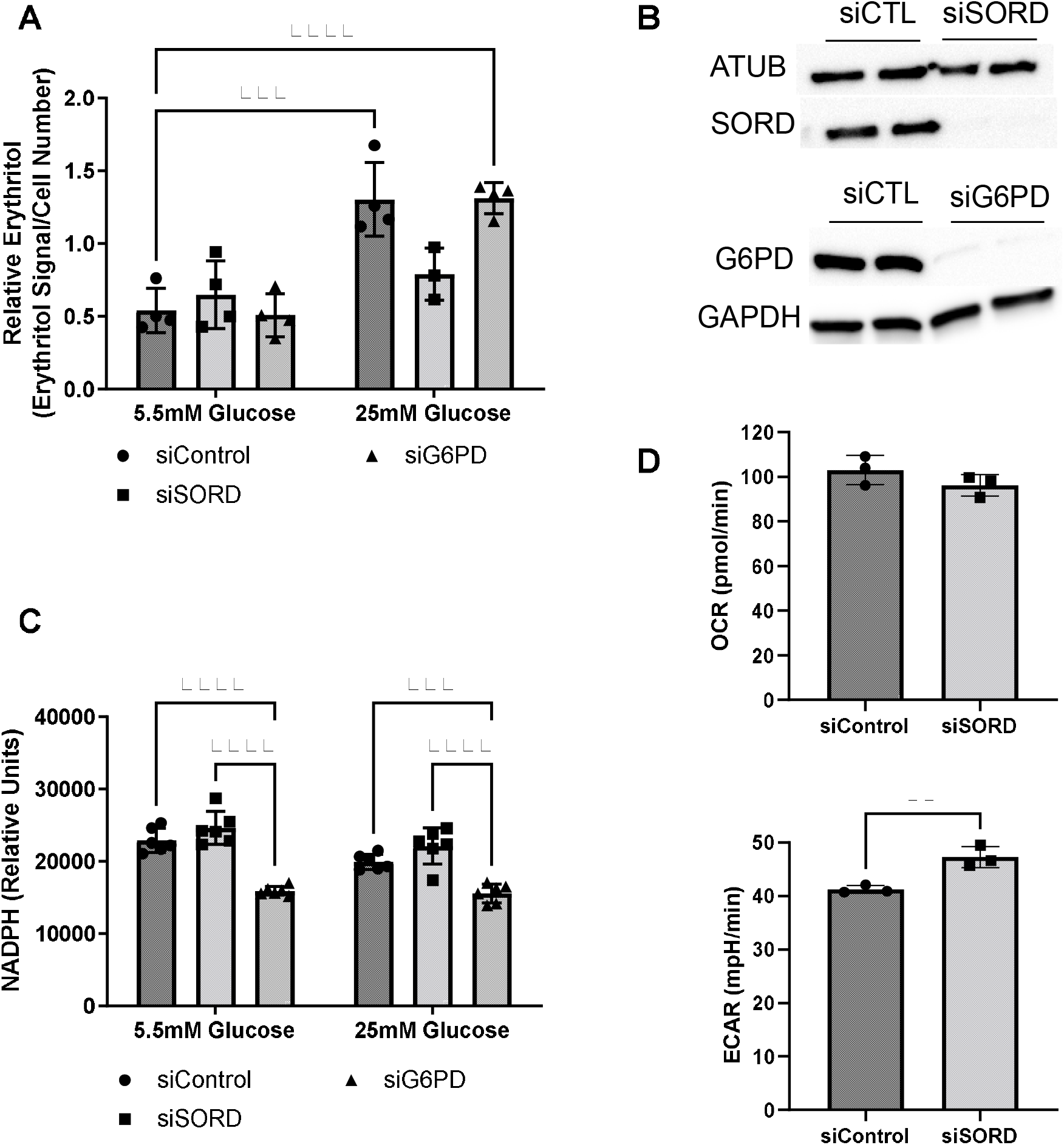
SORD and G6PD knockdown affect cellular metabolism. Knockdown of SORD and G6PD were performed by reverse transfection. (A) Intracellular erythritol (n=4), (B) protein levels of SORD and G6PD, and (C) intracellular NADPH were measured after 48hr treatment with 5.5mM or 25mM glucose (n=6). Erythritol was normalized to internal standard and cell number. (D) OCR and ECAR were measured in control or SORD knockdown cells following 48hrs in 10mM glucose media and normalized to cell number (n=3). Data is presented as mean ± SD, (A) and (C) were analyzed by two-way ANOVA followed by Sidak’s multiple comparisons test, (D) was analyzed with an unpaired t-test. **p<0.01, ***p<0.001, ****p<0.0001. ATUB, alpha tubulin; ECAR, extracellular acidification rate; GAPDH, glyceraldehyde 3-phosphate dehydrogenase; G6PD, glucose 6-phoshate dehydrogenase; SORD, sorbitol dehydrogenase.

Erythritol synthesis uses the coenzyme NADPH, which is produced by G6PD during the first steps of the PPP. We found that in both 5.5 mM and 25 mM glucose media, siG6PD significantly reduced intracellular NADPH compared to siControl and siSORD treated cells (Fig. 3C, p < 0.001). SORD knockdown did not significantly modify NADPH compared to control cells at either glucose concentration (Fig. 3C). Taken together, these findings suggest that erythritol synthesis is not affected by changes in intracellular NADPH.

We also assessed the effect of siSORD on cellular energy metabolism using real-time cell metabolic analysis (i.e. Agilent Seahorse technology). SORD knockdown significantly increased extracellular acidification rate (ECAR) (Fig. 3D, p < 0.01) and did not significantly change the oxygen consumption rate (OCR) (Fig. 3D). The increase in ECAR without significant changes to OCR likely indicates that when SORD expression is reduced, glycolysis and subsequent secretion of lactate are elevated.

### 3.3 Knockdown of the non-oxidative PPP enzyme transketolase reduces erythritol synthesis

The PPP consists of an oxidative phase, for which G6PD is rate-limiting, and a non-oxidative phase. The non-oxidative PPP, which provides erythrose (via erythrose-4-phosphate) for erythritol synthesis, requires the activities of transketolase (TKT) and transaldolase (TALDO) enzymes. Indeed, knockdown of TKT expression in cell cultured in high-glucose media significantly reduced intracellular erythritol levels (Fig. 4A, p < 0.01). This decrease in erythritol was of the same magnitude as siSORD (Fig. 4A, p < 0.01). Reduction of TALDO, however, did not significantly reduce intracellular erythritol compared to siControl cells (Fig. 4A). Successful knockdown of these enzymes was validated by western blot (Fig. 4B and 4C).

**Figure 4.**
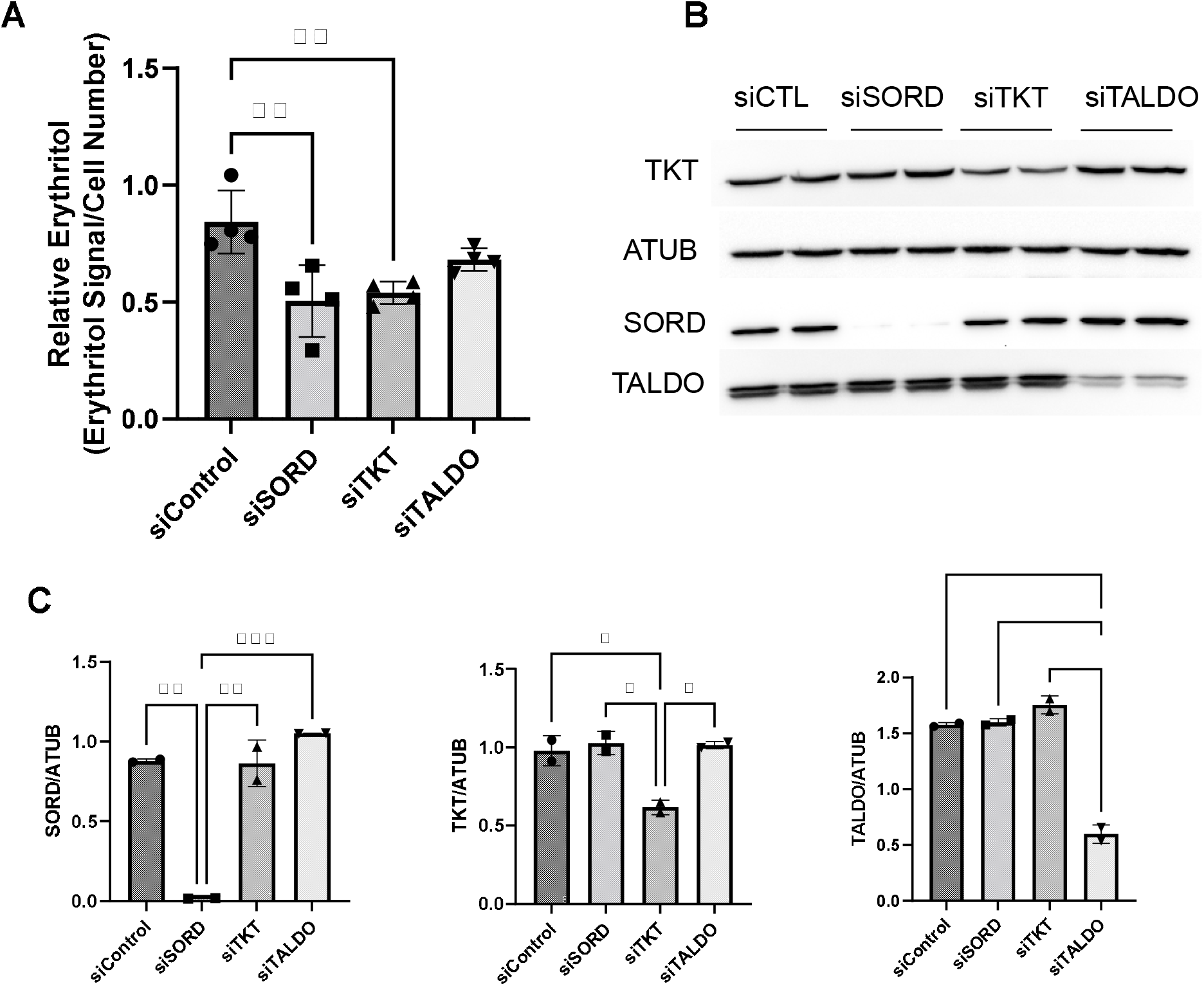
Knockdown of non-oxidative PPP enzymes SORD and TKT decrease intracellular erythritol levels. Knockdown of SORD, TKT, and TALDO were performed using reverse transfection. After 48hr treatment with 25mM glucose media, (A) intracellular erythritol (n=4) and (B) protein expression was measured. (C) Protein expression was quantified using Image J (n=2). Relative erythritol is normalized to internal standard and cell number. Data is shown as mean ± SD and was analyzed by ordinary one-way ANOVA and Tukey’s multiple comparisons test. *p<0.05, **p<0.01, ***p<0.001. ATUB, alpha tubulin; CTL, control; SORD, sorbitol dehydrogenase; TALDO, transaldolase; TKT, transketolase.

Given the effect of increasing glucose in media on erythritol synthesis, we assessed if high glucose also effected SORD and TKT expression. We found no significant difference SORD or TKT expression between 5.5 mM and 25 mM glucose-treated cells (Fig. 5A and 5B).

**Figure 5.**
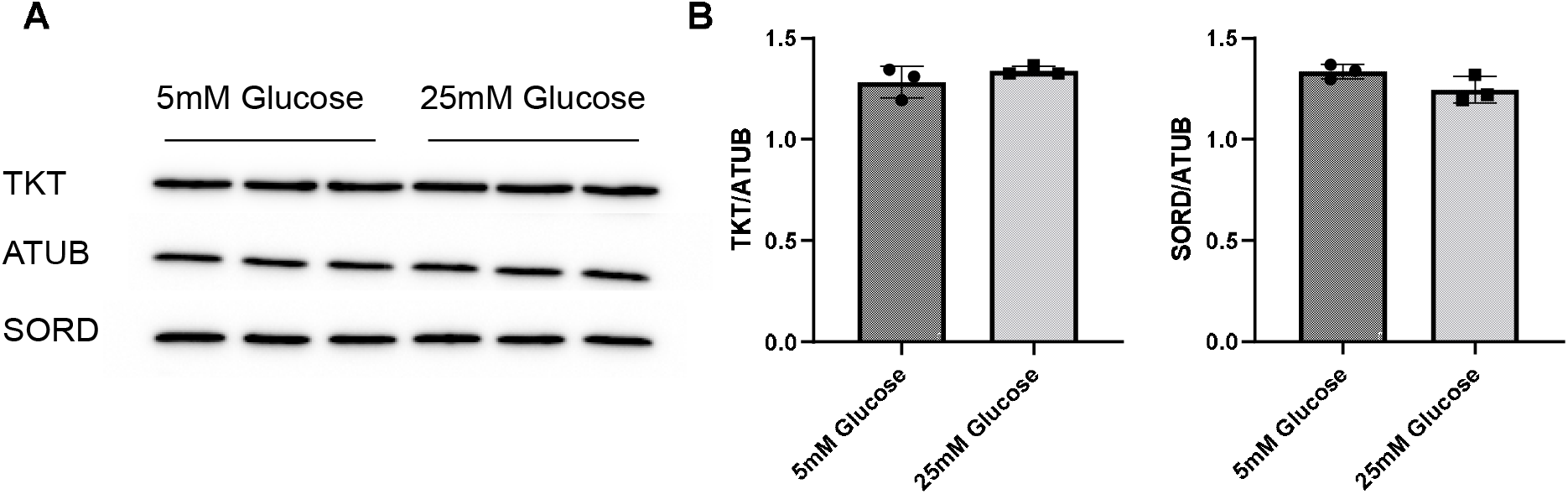
SORD and TKT protein expression do not increase in response to high glucose media. (A) Western blot of TKT and SORD expression after 48hr treatment in 5.5mM or 25mM glucose media and (B) quantification. Expression was quantified with ImageJ (n=3). ATUB, alpha tubulin; SORD, sorbitol dehydrogenase; TKT, transketolase.

### 3.4 Glucose-derived erythritol carbons originate from the PPP

We used position-specific [^13^C]-glucose tracing to determine if carbons for erythritol synthesis must first pass through the oxidative PPP, or can be derived directly from glycolysis through fructose-6-phosphate and glyceraldehyde-3-phosphate catalyzed by TKT and TALDO (Fig. 6A). 1-[^13^C]-glucose would be incorporated into erythritol through glycolysis, whereas 6-[^13^C]-glucose is incorporated through the PPP (Fig. 6A). We found that in cells treated with 1-[^13^C]-glucose, incorporation of labelled carbons into erythritol was 0% (M1 erythritol) (Fig. 6B). Contrastingly, treatment with 6-[^13^C]-glucose in siControl cells resulted in 60% M1 erythritol incorporation (Fig. 6C). Treatment with siSORD or siTKT also significantly reduced incorporation of 6-[^13^C]-glucose carbons into M1 erythritol (Fig. 6C, p < 0.0001 and p < 0.05). Consistent with previous findings, under high glucose conditions all erythritol carbons are derived from the oxidative PPP, but erythritol synthesis is limited by the enzymes SORD and TKT in the non-oxidative PPP [6,7].

**Figure 6.**
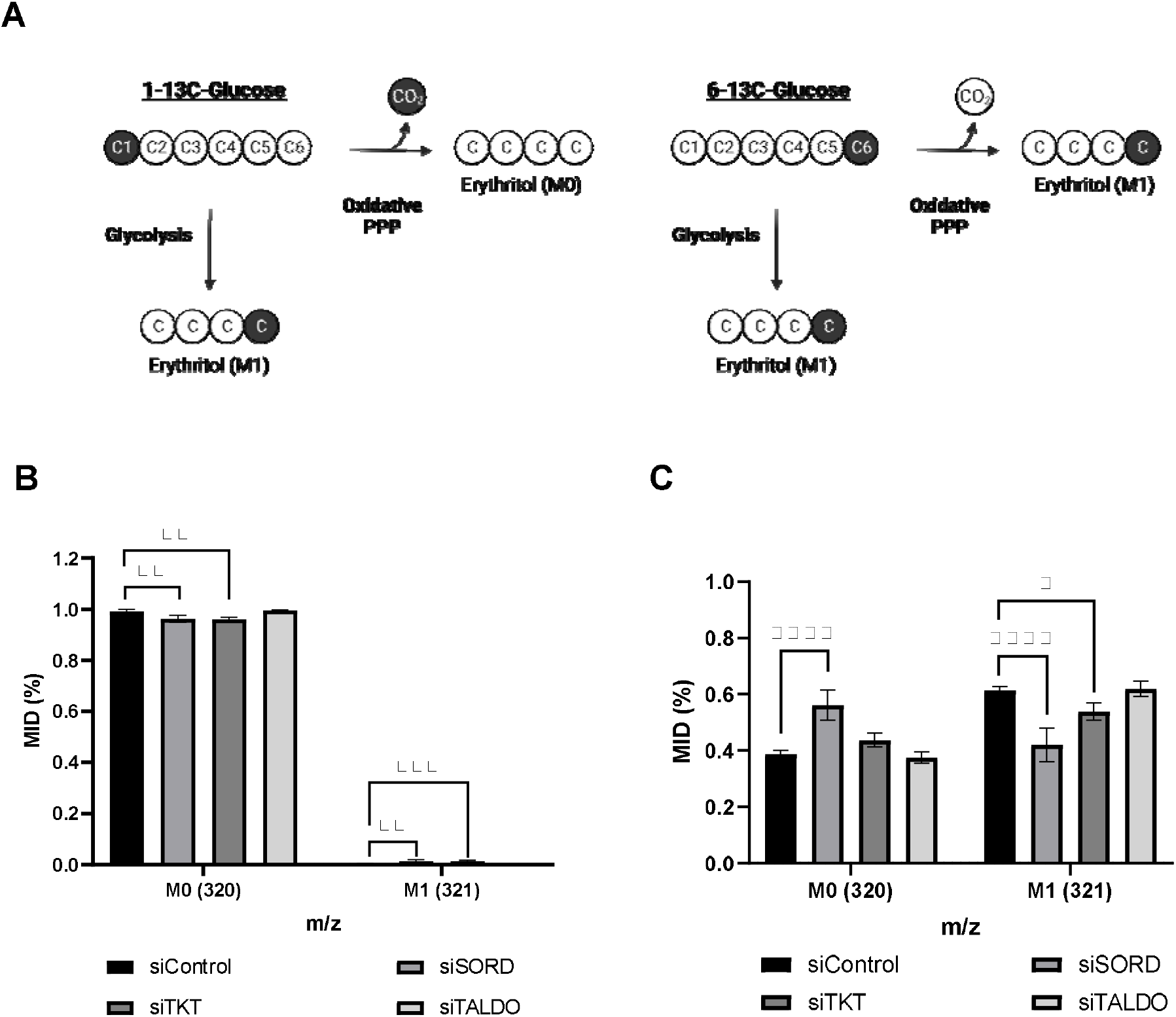
Erythritol carbons are derived from the oxidative PPP, not directly from glycolysis. (A) Representation of differential incorporation of ^13^C into erythritol from 1-[^13^C] or 6-[^13^C]-glucose. (B) MID of unlabeled (M0) and labelled (M1) erythritol after incubation with 25mM 1-[^13^C]-glucose. (C) MID of unlabeled (M0) and labelled (M1) erythritol after incubation with 25mM 6-[^13^C]-glucose. Data is shown as mean ± SD, n=4-5, and was analyzed by two-way ANOVA with Dunnett’s multiple comparisons test. *p<0.05, **p<0.01, ***p<0.001, ****p<0.0001. ATUB, alpha tubulin; MID, mass isotope distribution; SORD, sorbitol dehydrogenase; TALDO, transaldolase; TKT, transketolase.

### 3.5 Oxidative stress induces erythritol synthesis

We found that treatment with hydrogen peroxide for 8 hours significantly increased intracellular erythritol without significantly decreasing the percentage of live cells (Fig. 7A, p < 0.0001, and 6B). As expected, hydrogen peroxide also caused a significant increase in oxidized glutathione (Fig. 7C, p < 0.0001) and reduction in intracellular NADPH (Fig. 7D, p < 0.01). These findings suggest that erythritol synthesis is increased in response to oxidative stress and support the previous observation that decreased NADPH levels do not affect erythritol synthesis capacity (Fig. 3C).

**Figure 7.**
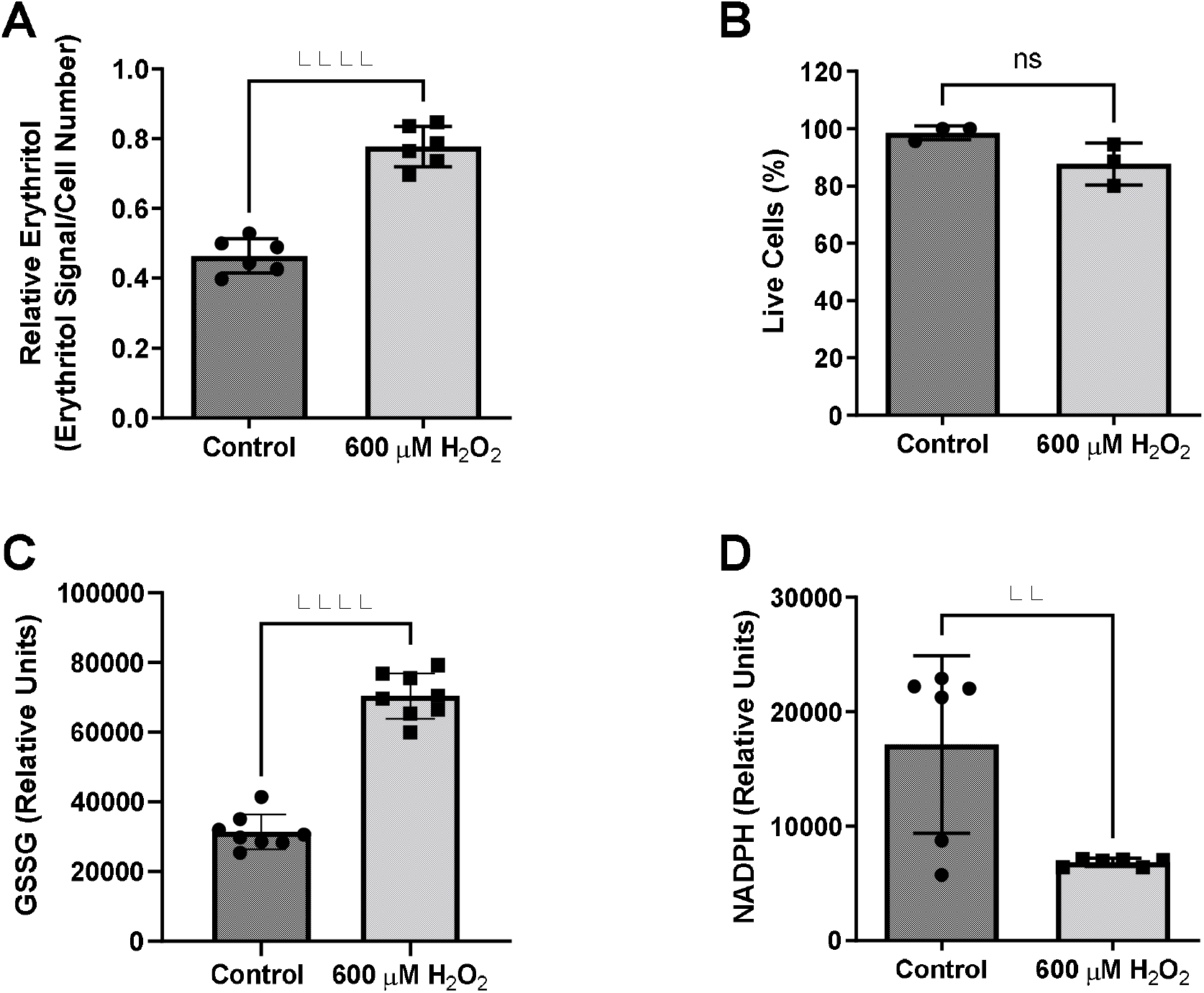
Intracellular erythritol and NADPH respond to hydrogen peroxide in culture medium. Cells were exposed to 600µM hydrogen peroxide or vehicle for 8 hr, after which (A) intracellular erythritol, (B) percentage of live cells, (C) oxidized glutathione (GSSG), and (D) NADPH were measured. Erythritol is normalized to internal standard and cell number. Data is presented as mean ± SD, n=3-6, analyzed by unpaired t-test. **p<0.01, ****p<0.0001. GSSG, oxidized glutathione; H_2_O_2_, hydrogen peroxide.

### 3.6 Keap1 mutations alter response to glucose and oxidative stress

We hypothesized that the increase in intracellular erythritol in response to high glucose and oxidative stress may be mediated by NRF2. To test this hypothesis, we utilized A549 cells that stably express either ectopic wildtype (WT) KEAP1 or the cysteine-to-serine mutations C273S and C151S [15,16]. The mutation C273S impairs the ability of KEAP1 to repress NRF2, resulting in constitutively active NRF2. C151S results in constitutive repression of NRF2 [16].

We confirmed that the A549 KEAP1 mutants differentially express KEAP1 and G6PD, a downstream target of NRF2 (Fig. 8A and 8B). Cells expressing WT, C151S, and C273S all significantly increase intracellular erythritol in 25mM glucose media (Fig. 9A, p < 0.001). The magnitude of this response, however, differed by KEAP1 status: C273S cells had 2 and 3-fold higher erythritol in 25mM glucose media compared to WT and C151S-expressing cells (Fig. 9A). This finding indicates that constitutively active NRF2 results in the most robust increase in erythritol synthesis from exposure to high-glucose in culture medium. We found a similar pattern in intracellular sorbitol. All cells responded to 25mM glucose by increasing sorbitol (Fig. 9B, p < 0.0001), but C273S cells accumulated 2-fold more sorbitol than cells with lower NRF2 expression.

**Figure 8.**
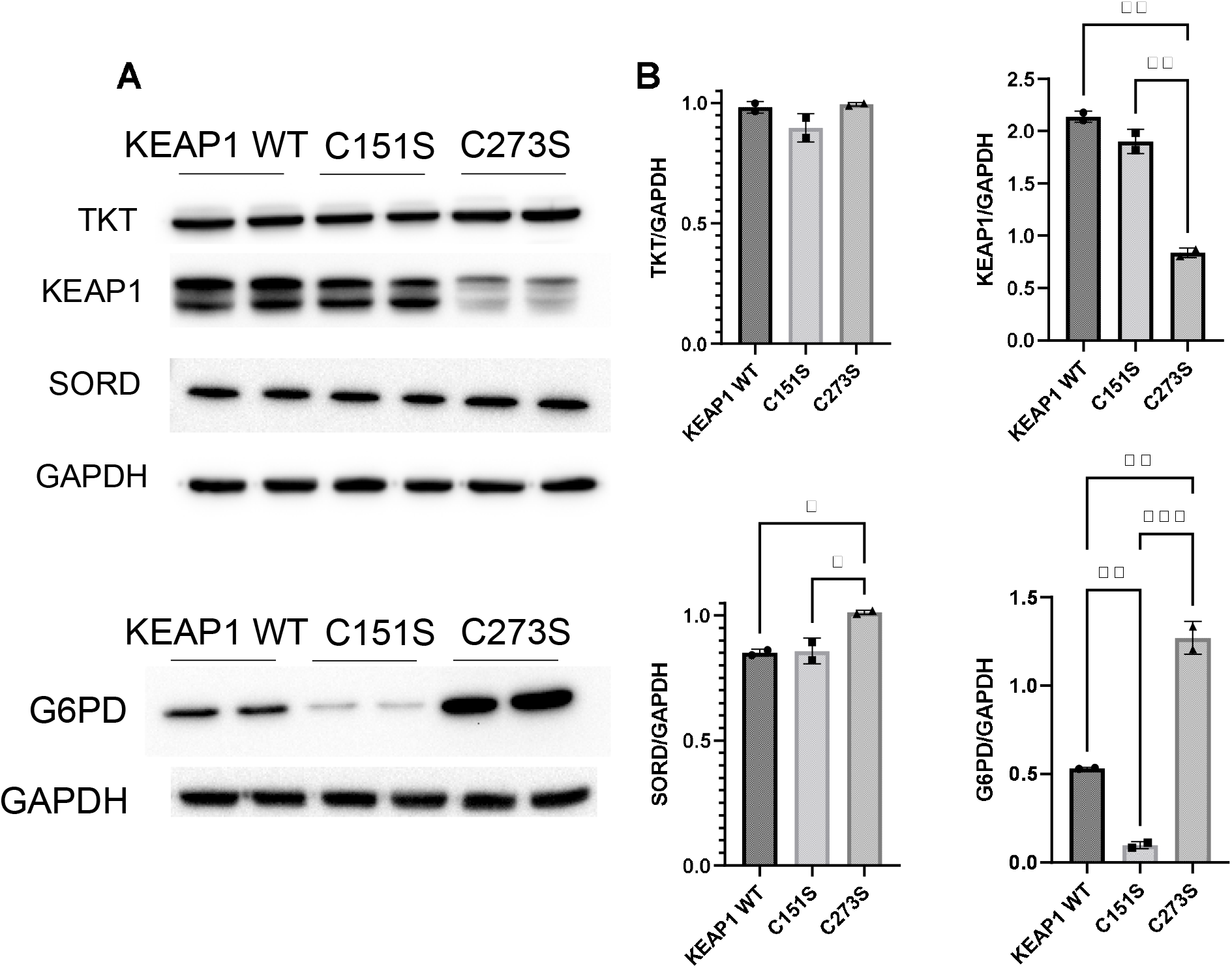
*KEAP1* variants have modified expression of proteins regulated by NRF2. (A) Western blots of TKT, KEAP1, SORD, and G6PD expression in KEAP1 mutant cells and (B) quantification of protein expression. Expression was quantified with ImageJ (n=3). G6PD, glucose-6-phosphate dehydrogenase; GAPDH, glyceraldehyde-3-phosphate dehydrogenase; KEAP1, Kelch Like ECH Associated Protein 1; SORD, sorbitol dehydrogenase; TKT, transketolase.

**Figure 9.**
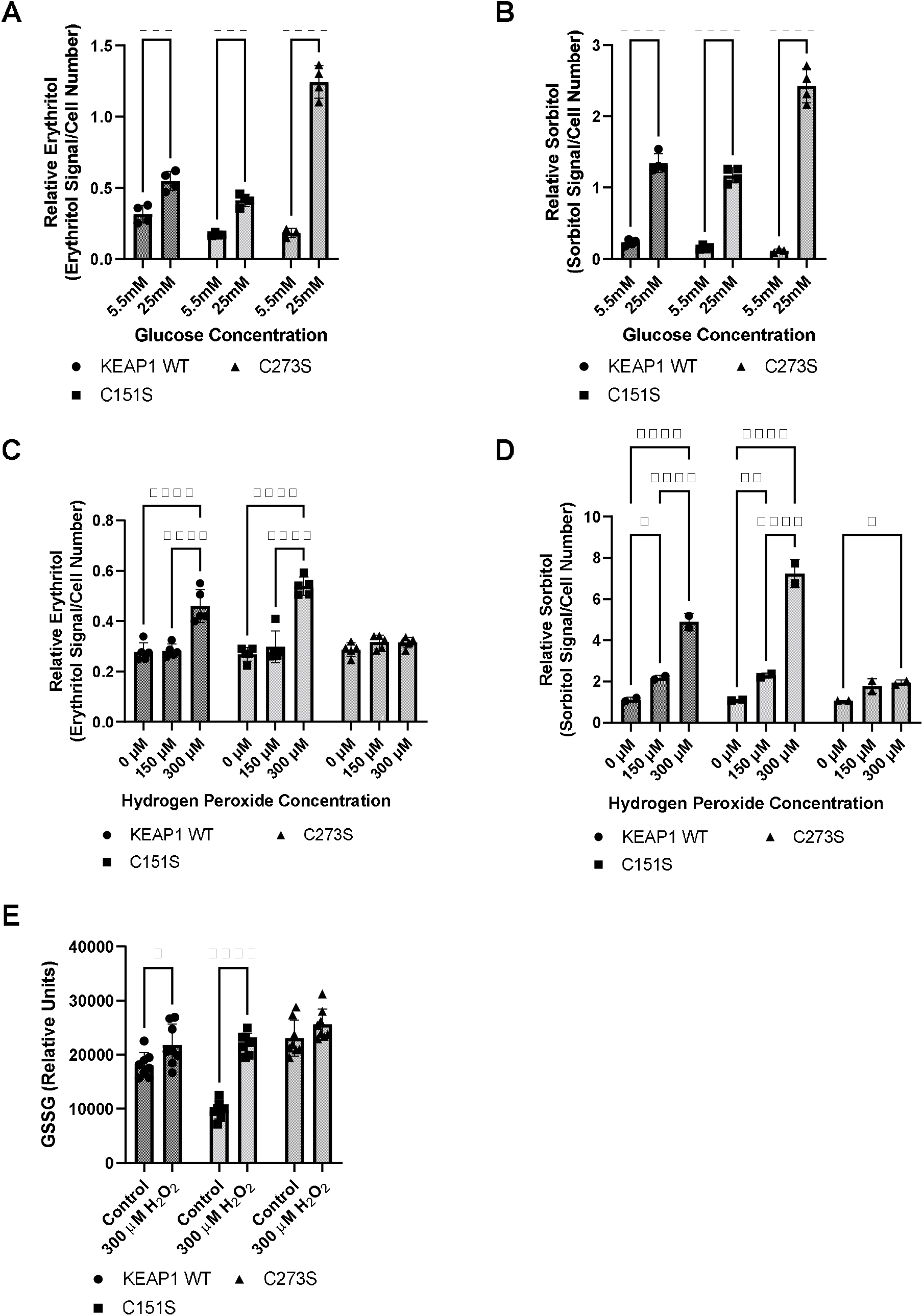
*KEAP1* variants modulate response to high glucose and oxidative stress. Intracellular erythritol (A) and sorbitol (B) in KEAP1 variant cells following 48 hr treatment with 5.5mM or 25mM glucose (n=4). Erythritol (C) and sorbitol (D) following exposure to 0-300µM hydrogen peroxide for 5 hrs (n=2-5). (E) Oxidized glutathione following 5 hr treatment with hydrogen peroxide. Erythritol is normalized to internal standard and cell number. Data is presented as mean ± SD and analyzed by two-way ANOVA followed by Sidak’s (A, B, E) and Tukey’s (C, D) multiple comparisons test. * p < 0.05, *** p < 0.001, **** p < 0.0001. WT-KEAP1 cells stably overexpress KEAP1. C273S cells have impaired ability of KEAP1 to repress NRF2, resulting in constitutively active NRF2. C151S cells exhibit constitutive repression of NRF2. GSSG, oxidized glutathione; H_2_O_2_, hydrogen peroxide.

Treatment with 300 µM hydrogen peroxide for 5 hours elevated erythritol in KEAP1 WT and C151S cells (Fig. 9C, p < 0.0001). There was no difference in C273S cells (Fig. 9C). Oxidized glutathione was also significantly elevated in KEAP1 WT and C151S, but not C273S cells (Fig. 9E, p < 0.05). Importantly, treatment with 600 µM hydrogen peroxide caused significant cell death in KEAP1 WT and C151S cells, indicating impaired response to oxidative stress in cells with reduced NRF2 activity. This finding is supported by the relative resistance of the C273S cells, which have constitutively active NRF2. Intracellular sorbitol also increased in response to 300 µM hydrogen peroxide in all 3 cell types, which indicates an accumulation of intracellular glucose (Fig. 9D, p < 0.05) and possible inhibition of glycolysis under these conditions.

## 4. Discussion

Previous literature has identified that, in mammals, erythritol is synthesized from glucose through the pentose phosphate pathway [6]. Erythritol synthesis from erythrose can be catalyzed by SORD and ADH1 using NADPH as a cofactor [7]. The precursor of erythrose is erythrose-4-phosphate, which is a product of the non-oxidative PPP. The regulation of erythritol synthesis and its role in metabolic homeostasis, however, is poorly understood [17]. Here, we demonstrate that erythritol synthesis is modulated both by glucose availability and oxidative stress. We observed that intracellular erythritol increased dose-dependently with increasing glucose in culture media (Fig. 1A). Interestingly, we also demonstrated that in the absence of glucose, erythritol synthesis is still elevated by exposure to 25 mM fructose compared to 6.25 mM glucose (Fig. 1A). This elevation is modest, however, when compared to the response to 25 mM glucose media. In A549 cells, fructose is converted to glycolytic intermediates that are primarily used for fatty acid synthesis [18,19]. Fatty acid oxidation utilizes NADPH, thus increasing the demand for NADPH regeneration in the oxidative pentose phosphate pathway [19]. Fructose, then, may promote erythritol synthesis directly through increased glycolytic intermediates or through increased PPP flux.

In addition to increased nutrient availability, excess carbohydrates are a source of osmotic stress in cell culture. In yeast, erythritol synthesis can be induced by osmotic stress [20]. In humans, the 6-carbon polyol sorbitol is also known to be elevated during osmotic stress [14,21]. To evaluate if erythritol synthesis is an additional mechanism of osmoregulation, we treated cells with mannitol. Mannitol is strong osmolyte that undergoes little metabolism in humans [22]. Interestingly, mannitol treatment did not affect intracellular erythritol, whereas sorbitol was significantly elevated (Fig 1B, 1C). This may be explained by differing capacities for diffusion across the plasma membrane. Erythritol more readily diffuses across cell membranes than the larger polyol sorbitol [22]. Sorbitol, therefore, is likely a more effective endogenous osmolyte to combat hyperosmotic stress than is erythritol.

Yeast also use glycerol as a substrate for erythritol synthesis by converting glycerol to glyceraldehyde-3-phosphate, a precursor of erythrose-4-phosphate in the non-oxidative PPP [20]. Based on the finding that fructose, another direct precursor of erythrose-4-phosphate, elevated erythritol, we expected that glycerol treatment would produce similar results in human cells. In further support that erythritol does not respond to osmotic stress in human cells, 0.5 mM glycerol significantly increased intracellular sorbitol, but did not impact intracellular erythritol (Fig 1D, 1E). Collectively, these data support that erythritol is produced in response to carbohydrate abundance and not in response to osmotic stress.

A549 cells are derived from cancerous tissue, which is known for high PPP activity [23,24]. We aimed to evaluate if non-cancerous cells exhibit the same increase in erythritol in response to high glucose conditions. We chose to use HK-2 cells based on our previous work showing that the kidney contains relatively high endogenous erythritol in mice [12]. Indeed, we found that HK-2 cells also respond to excess glucose with an increase in intracellular erythritol (Fig 2), consistent with previous observations of increased PPP flux in HK-2 cells cultured in hyperglycemic hypoxic conditions [25]. This demonstrates that glucose availability can also regulate synthesis in non-cancerous erythritol-producing cells.

Our work validated the previous finding that SORD knockdown significantly decreases erythritol synthesis [7], as previous experiments were conducted in media containing 25 mM glucose [7]. This is the first study to report that the effect of SORD knockdown on intracellular erythritol is dependent on glucose level. In basal glucose media (5.5 mM), siSORD does not significantly decrease intracellular erythritol whereas in high glucose media (25 mM glucose), siSORD caused a 40% reduction (Fig 3A). This further supports the primary role of glucose availability in the synthesis of erythritol.

G6PD is the rate-limiting enzyme of the PPP. We expected, then, that knockdown of G6PD would result in reduced intracellular erythritol in high-glucose media. Interestingly, G6PD knockdown did not blunt the glucose-induced increase in erythritol (Fig. 3A). One explanation for this is the paradoxical finding by Zhao et al. that in A549 cells, the constitutive activation of NRF2 results in both high expression of oxidative PPP enzymes and reduced dependence on the oxidative PPP for cell growth [15]. Another study in melanoma cells, which also have high PPP activity, found that when G6PD function was impaired, there was no reduction in erythrose-4-phosphate levels [26]. These studies demonstrate that cancer cells are resilient to G6PD inhibition and may continue to fuel the non-oxidative PPP through alternative methods.

We next evaluated the impact of knocking down the downstream non-oxidative PPP enzymes TKT and TALDO. We found that knocking down *TKT*, but not *TALDO*, reduced erythritol in 25 mM glucose media by a similar magnitude as *SORD* knockdown (Fig. 4A). This is consistent with historic reports that *TALDO* deficiency resulted in accumulation, rather than depletion, of erythritol and other polyols in plasma and urine [27,28]. Our findings indicate that both *SORD* and *TKT* expression are essential for the synthesis of erythritol.

TKT participates in two reversible sugar conversions in the non-oxidative PPP. The TKT-catalyzed reaction erythrose-4-phosphate + xylulose-5-phosphate ? fructose-6-phosphate + glyceraldehyde-3-phosphate is a bridge by which carbons can be passed between glycolysis and the PPP. Because siG6PD did not limit erythritol synthesis, but siTKT did, we aimed to understand if erythritol synthesis is being supported by carbons directly from glycolytic intermediates when glucose availability is high. We found using position-specific [^13^C]-glucose tracing that when G6PD expression is not altered, glucose passes through the oxidative PPP before incorporation into erythritol (Fig. 6 A-C). This is in agreement with the previous finding that in A549 cells, all erythritol was derived from glucose passed through the oxidative PPP [6].

Flux through the PPP is a key defense mechanism to combat oxidative stress, primarily through generation of reducing equivalents as NADPH. We hypothesized that oxidative stress, therefore, would also elevate synthesis of erythritol. As expected, we found that intracellular erythritol is elevated in A549 cells exposed to hydrogen peroxide (Fig. 7A). Interestingly, erythritol synthesis capacity is not associated with intracellular NADPH levels—in fact, erythritol was elevated with hydrogen peroxide treatment when NADPH was depleted (Fig. 7D). Similarly, G6PD knockdown depleted intracellular NADPH, but did not affect erythritol levels (Fig. 3C). Together, these findings suggest that flux of glucose through G6PD and intracellular NADPH are sufficient, even when NADPH is reduced, to support erythritol synthesis.

We further explored the relationship between oxidative stress and erythritol utilizing A549 *KEAP1* mutant cells with altered activity of NRF2 [15,16]. NRF2 directs glucose flux through the PPP by modifying enzyme expression [29,30]. We found that constitutively active NRF2 intensified glucose-induced erythritol synthesis, resulting in even higher erythritol levels than cells with normal or impaired NRF2 (Fig. 9A). Notably, cells with constitutively active NRF2 did not have higher intracellular erythritol at baseline. This suggests that the co-occurrence of hyperglycemia and oxidative stress may be a key factor in elevating erythritol synthesis.

We also found that impairing NRF2 did not eliminate erythritol synthesis during oxidative stress but did lower the threshold for this response. In parental A549 cells, 600 µM hydrogen peroxide induced oxidative stress and elevated intracellular erythritol (Fig. 7A). This dose was highly cytotoxic to cells with ectopic WT KEAP1 or the NRF2-repressing C151S mutation. WT KEAP1 and C151S cells increased erythritol synthesis following treatment with half the dose, 300 µM hydrogen peroxide (Fig. 9C). Erythritol synthesis during oxidative stress may be due both NRF2-dependent and NRF2-independent mechanisms, as oxidative stress inhibits the activity of several glycolytic enzymes through mechanisms that are not dependent on NRF2, promoting the accumulation of glycolytic intermediates [23,24]. We also observed significant accumulation of sorbitol in KEAP1 WT and C151S cells under oxidative stress (Fig. 9D). This supports that intracellular glucose is elevated, likely due to the inhibition of glycolytic enzymes, which promotes alternative pathways (i.e. sorbitol and erythritol synthesis through the polyol and PPP, respectively). Overall, our findings in KEAP1 mutant cells further indicate that elevated erythritol synthesis is a marker of glucose flux through the PPP.

In humans, elevated circulating erythritol is a predictive biomarker of cardiometabolic diseases [17]. Our findings provide a connection between erythritol regulation and hallmarks of cardiometabolic disease: hyperglycemia and oxidative stress [31,32]. We demonstrated that erythritol is elevated by both hyperglycemia and oxidative stress, and that these effects can compound to further promote erythritol synthesis. Further research characterizing whole-body erythritol homeostasis may provide a powerful tool for detecting early cardiometabolic dysfunction.

## Acknowledgements

The authors are grateful to Christian Metallo for the generous gift of the A549 KEAP1 over-expression cell lines (WT and mutants).

## Author Contributions

**Semira R. Ortiz:** Conceptualization, Methodology, Investigation, Visualization, Writing – Original draft preparation. **Alexander Heinz:** Methodology and Software. **Karsten Hiller**: Methodology. **Martha S. Field**: Conceptualization, Supervision, Writing – Review and Editing.

