## supplemental table 1 for "Erythritol synthesis in human cells is elevated in response to oxidative stress and regulated by the non-oxidative pentose phosphate pathway"

**Supplementary Tables**

**Table S1**

| **Gene** | **Catalog Number** | **Sequences** |
| --- | --- | --- |
| Non-targeting pool | D-001810-10-05 | UGGUUUACAUGUCGACUAA, UGGUUUACAUGUUGUGUGA, UGGUUUACAUGUUUUCUGA, UGGUUUACAUGUUUUCCUA |
| SORD | L-008323-00-0005 | CAGAAUCCCUGAUGUUAAU, GAUCAUCGGUAAAGCACCU, GAAAUGUCAUGUGAGGUUA, GAUACAAUCUGGGAUAGUU |
| G6PD | L-008181-02-0005 | ACAGAUACAAGAACGUGAA, CCGUGUACACCAAGAUGAU, CAGAUAGGCUGGAACCGCA, AUUCACGAGUCCUGCAUGA |
| TKT | L-004734-00-0005 | GGAACUAGCCGCCAAUACA, CCGUGGAGGACCAUUAUUA, GCAGUUAACCGGGUACCAA, GAUAAGGAGUCUUGGCAUG |
| TALDO1 | L-008996-00-0005 | GCAAACACCGACAAGAAAU, UCACAAGAGGACCAGAUUA, CCGAGUAUCCACAGAAGUA, ACAAGAAGUUUAGCUACAA |
